# Investigation of singlet oxygen sensitive genes in the cyanobacterium *Synechocystis* PCC 6803

**DOI:** 10.1101/2023.09.22.558817

**Authors:** Gábor Patyi, Barbara Hódi, Ivy Mallick, Gergely Maróti, Péter B. Kós, Imre Vass

## Abstract

Singlet oxygen (^1^O_2_) is an important reactive oxygen species whose formation by the type-II, light-dependent, photodynamic reaction is inevitable during photosynthetic processes. In the last decades, the recognition that ^1^O_2_ is not only a damaging agent, but can also affect gene expression and participates in signal transduction pathways has received increasing attention. However, contrary to several other taxa, ^1^O_2_-specific genes have not been identified in the important cyanobacterial model organism *Synechocystis* PCC 6803. By using global transcript analysis we have identified a large set of ^1^O_2_-responsive *Synechocystis* genes, whose transcript levels were either enhanced or repressed in the presence of ^1^O_2_. Characteristic ^1^O_2_ responses were observed in several light-inducible genes of *Synechocystis*, especially in the *hli* (or *scp*) family encoding HLIP/SCP proteins involved in photoprotection. Other important ^1^O_2_-induced genes include components of the Photosystem II repair machinery (*psbA2* and *ftsH2*, *ftsH3*), iron homeostasis genes *isiA* and *idiA*, the Group-2 sigma factor *sig*D, some components of the transcriptomes induced by salt-, hyperosmotic and cold-stress, as well as several genes of unknown function. One of the most pronounced ^1^O_2_-induced upregulation was observed for the *hliB* gene, whose deletion provided tolerance against ^1^O_2_-mediated light damage. A bioreporter *Synechocystis* strain was created by fusing the *hliB* promoter to the bacterial luciferase (lux), which showed its utility for continuous monitoring of ^1^O_2_ concentrations inside the cell.

## Introduction

Singlet oxygen (^1^O_2_) is an extremely important reactive oxygen species (ROS) with a high oxidative capacity. It can participate in diverse intracellular processes, damaging macromolecules, oxidizing proteins, fatty acids, and nucleotides (Krieger-Liszkay, 2005)(Di Mascio *et al*., 2019; Pospíšil *et al*., 2022), inducing adverse effects on cells. One of its most significant physiological effect in the photosynthetic apparatus is the damage of the structure and function of the Photosystem II (PSII) reaction centre under high light exposure (Okada *et al*., 1996; Mizusawa *et al*., 2003; Krieger-Liszkay, 2005; Krieger-Liszkay *et al*., 2008; Rehman *et al*., 2013; Bashir *et al*., 2021).

The reactive nature of ^1^O_2_ not only causes damage to various cell constituents, but can also induce changes in gene expression. In the green alga *Chlamydomonas reinhardtii* and higher plants it has been demonstrated that ^1^O_2_ acts as a signaling molecule that transmits information from chloroplasts to the nucleus, regulating the expression of nuclear genes (Fischer *et al*., 2007; Wang *et al*., 2016; Dmitrieva *et al*., 2020). ^1^O_2_-mediated activation of genes involved in the molecular defense response against photooxidative stress has been reported in various organisms (Leisinger *et al*., 2001; Op Den Camp *et al*., 2003; Berghoff *et al*., 2011). Leisinger et al. showed that in the presence of the external photosensitizer Rose bengal (RB) the glutathione peroxidase (*gpxh*) homologous gene of *Chlamydomonas* is transcriptionally activated by ^1^O_2_, whereas *gpxh* mRNA levels are only weakly increased by O_2_^•−^ or peroxide (Leisinger *et al*., 2001). Op den Camp et al. used the *Arabidopsis* flu mutant to demonstrate that ^1^O_2_ formed as a result of protochlorophyll accumulation rapidly activated many (70) genes (Op Den Camp *et al*., 2003). In contrast, other ROS, such as O_2_^•−^, did not affect the expression of these genes during the early stress response. Among photosynthetic prokaryotes gene level responses to ^1^O_2_ have been investigated only in the case of the phototrophic alpha-proteobacterium *Cereibacter sphaeroides* (old name: *Rhodobacter sphaeroides*) (Glaeser *et al*., 2011). Information on this important topic in the case of the widely used model cyanobacterium *Synechocystis* PCC 6803 is completely missing.

Investigation of signaling processes mediated by ^1^O_2_ inside cells requires methods that are suitable for ^1^O_2_ detection in the intracellular environment. However, the available methods, such as EPR spin trapping by TEMP (Hideg *et al*., 1994; Fufezan *et al*., 2007) or TEMPD-HCl (Leisinger *et al*., 2001; Ferretti *et al*., 2018), direct 1270 nm luminescence detection (Tomo *et al*., 2012), fluorescent spin traps, DanePy (Hideg *et al*., 2007) and SOSG (Flors *et al*., 2006; Bashir *et al*., 2021) are not suitable for continuous detection of intracellular ^1^O_2_ levels. The use of cyanobacterial bioreporters can provide a solution for this problem (Patyi *et al*., 2021), for which the identification of specific ^1^O_2_-responsive genes is indispensable.

In the present work we identified genes in *Synechocystis* whose expression was specifically upregulated or suppressed in the presence of ^1^O_2_ generated from either endogenous or exogenous sources. One of the most promising ^1^O_2_-specific genes is the high light inducible *hliB* whose expression level is upregulated ca. 50-fold by ^1^O_2_. Deletion of this gene enhanced ^1^O_2_-dependent light sensitivity of *Synechocystis*, demonstrating the involvement of the HliB protein in protection against ^1^O_2_-mediated photodamage. By using a fusion of the *hliB* promoter and the bacterial luciferase gene we created a ^1^O_2_ bioreporter construct, which allows detection of ^1^O_2_ production inside the cyanobacterial cells.

## Materials and Methods

### Strains, growth conditions

*Synechocystis* PPC 6803 (*Synecocystis*) cells were grown photoautotrophically in the presence of 3% CO_2_, 40 μmol photon m^−2^ s^−1^ white light intensity and 30°C in BG-11 (Rippka *et al*., 1979) as described earlier (Patyi *et al*., 2021).

*Escherichia coli* strain *DH5*α, used for routine DNA manipulations (Kirtania *et al*., 2019) and constructions, was grown in Luria broth (LB) medium at 37°C (J. Sambrook, D.W. Russell, 2001).

### Measurement of growth

The differences in growth rates caused by ^1^O_2_, produced by either 0.5 µM Methylene Blue (MB) or Rose Bengal (RB), were assessed by following the optical density at 680 nm for 4 days in a Photo Multi Cultivator MC-1000 (Photon Systems Instrument) with automatic OD measurements in every hour.

### Gene expression studies

*Synechocystis* cDNA libraries were prepared for whole transcriptome sequencing from 0.5 µg of total RNA using the NEBNext rRNA depletion Kit for Bacteria #E7850, #E7860 (Biolabs). Paired end sequencing was performed using the Illumina NextSeq platform with the NextSeq 500/550 High Output Kit v2.5 (75 Cycles) resulting in 306611978 reads altogether (10M reads per sample). Trimming, quality clipping, gene identification and downstream calculations were carried out in CLC Genomics Workbench 20.0 (QIAGEN). The *Synechocystis* genome sequence NC_000911 downloaded from NCBI was used as template for mapping the reads to genes. Gene abundances were calculated in RPKM (Reads Per Kilobase Million). For comparative study, *Log_2_ fold changes* and *p* values were calculated as well as *Max group means* were established for each gene. (For each group in the statistical comparison, the average RPKM is calculated. *Max group means* is the maximum of the average RPKM’s.)

To verify transcript abundances, the same total RNA samples were used as for total transcriptome sequencing. cDNA was synthesized using the RevertAid RT Kit (ThermoFisher Scientific) and used as template in quantitative PCR (qPCR) using 5X HOT FIREPol EvaGreen qPCR Mix Plus (Solis BioDyne) in the CFX384 Touch Real-Time PCR Detection System. The ΔΔC_T_ method was used for calculating changes of gene expression using *rrn16S* as internal control.

### Construction of the ΔhliB strain

We constructed a *ΔhliB* mutant by replacing the *hliB* (*ssr2595*) gene with a spectinomycin cassette (Omega casette). For creating the insert of a pUC19 vector, the spectinomycin cassette HindIII/BamHI fragment from the vector pND6LuxAB (Peca *et al*., 2008) was amplified and ligated between two 0.5-kb-long genomic fragments surrounding the *hliB gene.* The plasmid was amplified in *E. coli* and transformed to *Synechocystis* via natural transformation. The mutant formed by double crossover was grown on selective BG-11 plates containing 50 μg mL^−1^ spectinomycin.

### Construction of the hliBLux bioreporter strain

We used the pILA promoter probe vector (Kunert *et al*., 2000) utilizing the *LuxAB* luminescence reporter system. The insert of the *hliB* (*ssr2595*) promoter region was amplified by PCR using chromosomal DNA of WT *Synechocystis* as a template and the appropriate primer pair with restriction sites (Table 1).

**Table 1.**
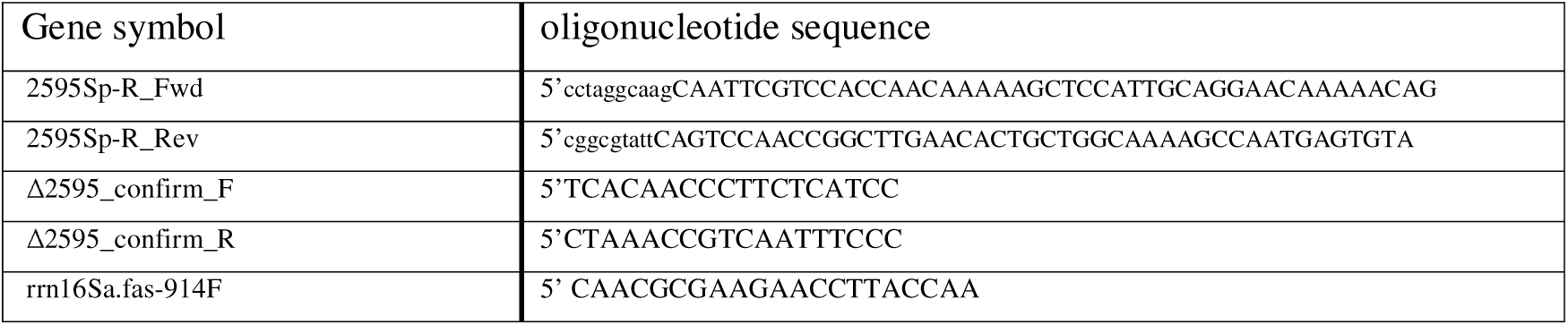
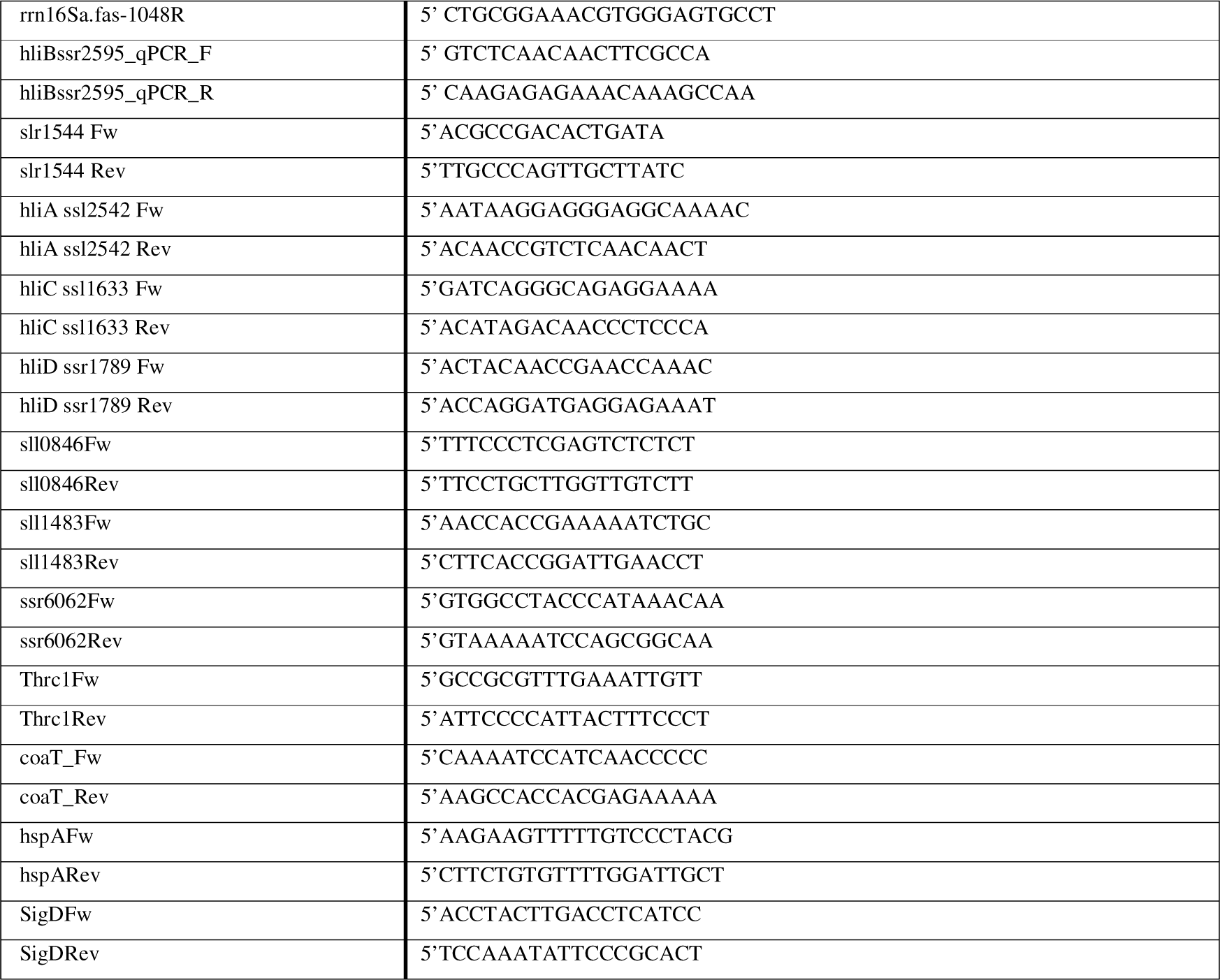
Oligonucleotides used in this work.

The promoter fragments were inserted into the unique PstI-KpnI site of pILA. *E. coli* transformants were grown on kanamycin selective LB plates. The pILA*hliB* plasmids were amplified and isolated from *E. coli*. The wild-type *Synechocystis* strain was transformed and the clones were selected with kanamycin selection as described above.

### Bioluminescence assay

^1^O_2_ treatments were carried out in 10 mL batches of *hliB*Lux *Synechocystis* culture in BG-11 medium at 30°C under LL and HL conditions. For the generation of exogenous ^1^O_2_ we used either MB or RB photosensitizer dyes in 0.5 µM concentration. When indicated, 5 mM histidine (His) was used as ^1^O_2_ scavenger (Rehman *et al*., 2013). Aliquots were withdrawn in every 15 minutes for luminescence measurements performed using the previously described protocol (Patyi *et al*., 2021).

### Data analysis and representation

The results were evaluated using the Microsoft Office Excel 2016 program and the OriginPro 2021b data analysis program. The *in silico* design of the molecular biology work and the analysis of our whole transcriptome data were performed using the CLC Genomics Workbench 20.0 (QIAGEN). Figures 4A and 5A were created using the UpSet online intersecting sets visualization program (Lex *et al*., 2014).

## Results

### Global transcript analysis of gene expression changes induced by ^1^O_2_

To identify genes, whose expression is affected by ^1^O_2_, we exposed *Synechocystis* cells to ^1^O_2_ generated either by HL illumination in the absence of any addition (endogenous ^1^O_2_), or by illumination in the presence of MB or RB (exogenous ^1^O_2_). This illumination protocol allowed us to monitor changes in the expression of genes that are sensitive to ^1^O_2_ generated via pigment molecules of the photosynthetic apparatus, as well as by the added photosensitizers. The application of His as a ^1^O_2_ scavenger allowed the confirmation of the ^1^O_2_ specificity of the genes.

In the first step of transcript analysis we compared the transcript abundance of samples kept in low light with that of samples exposed to high light (LL vs HL). A strong effect of HL treatment on gene expression was visible for about 500 genes showing changes in expression in the form of either induction (log_2_FC > 0.9) or repression (log_2_FC < −0.9) (Fig. 3A).

**Figure 1.**
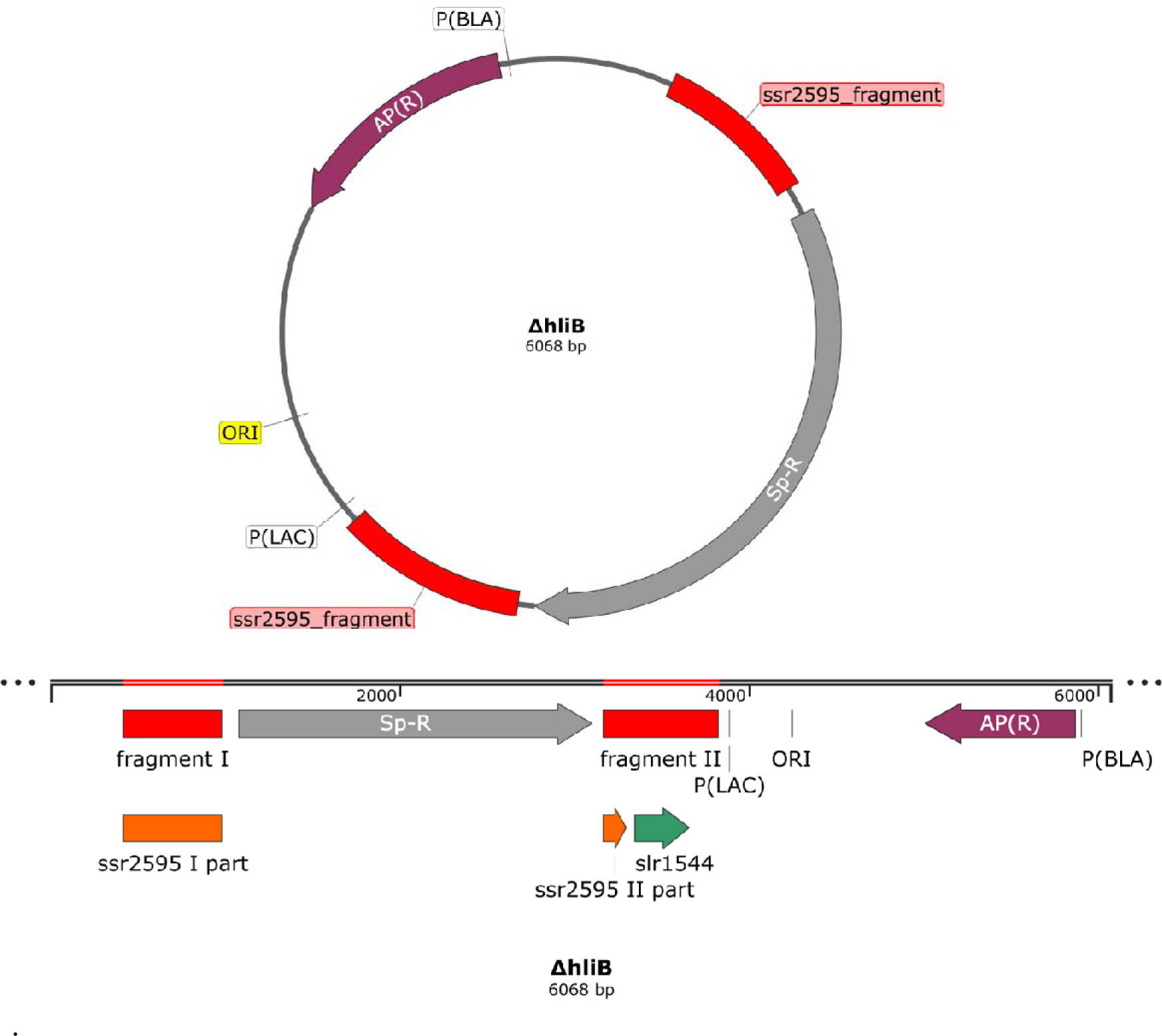
Schematic representation of the mutagenic plasmid used for construction of the *ΔhliB* strain. The figure shows the plasmid map of the pUC19 vector with the ssr2595-Spe^R^-ssr2595 fragment.

**Figure 2.**
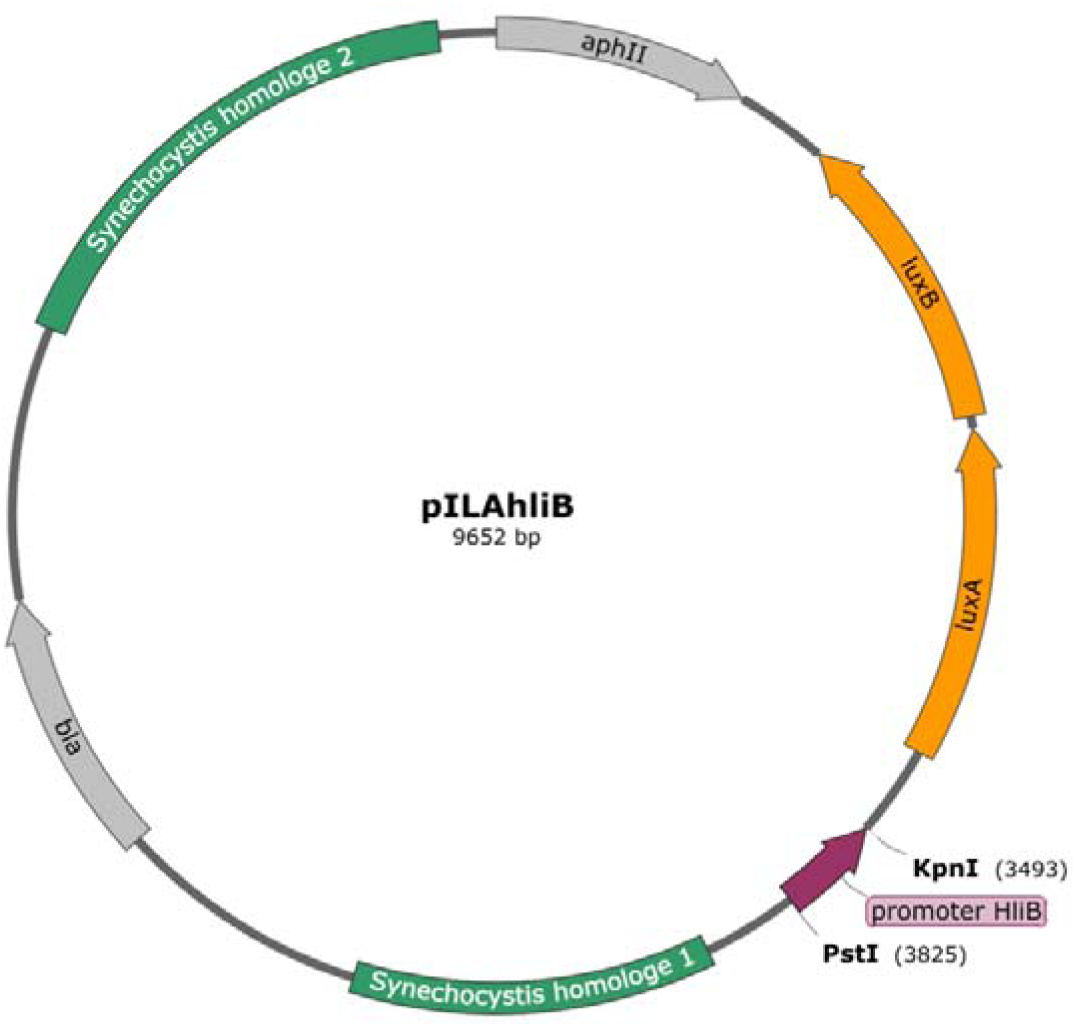
Schematic figure of the pILA promoter probe vector with the insert of *hliB* promoter. We used the appropriate primers (Table 1) for the amplification of fragments from the WT *Synechocystis* genome and applied KpnI-PstI digestion. *luxA* and *luxB* genes code for the luciferase reporter proteins, which are required for the emission of detectable bioluminescence. *aphII* and *bla* refer to genes conferring resistance to ampicillin and kanamycin, respectively.

**Figure 3.**
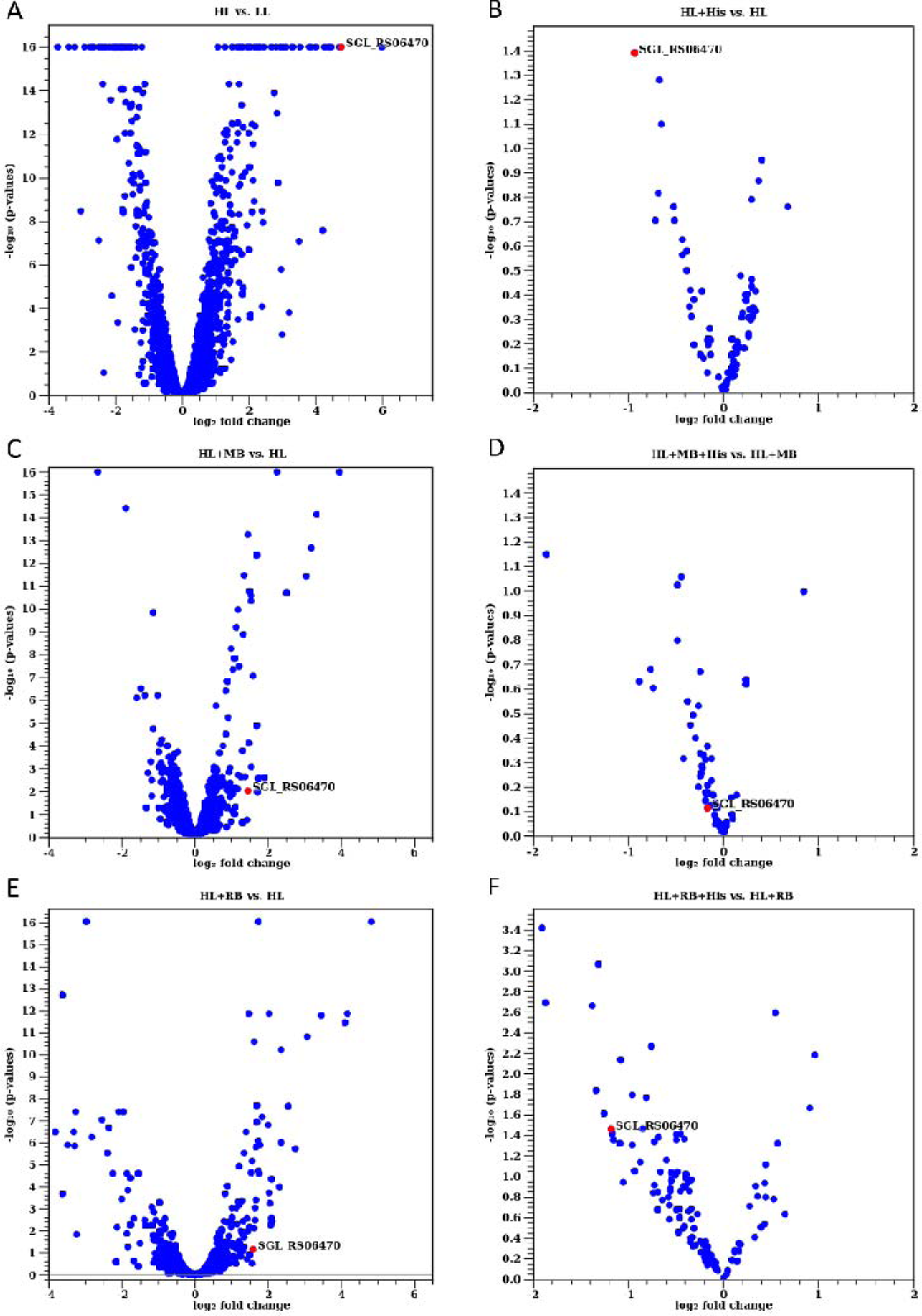
Volcano plots of the global transcript analysis of wild-type *Synechocystis* cells. The cyanobacterial cells were exposed to low (LL: 30 μmol photon m^−2^ s^−1^) or high (HL: 200 μmol photon m^−2^ s^−1^) light intensity in the absence or presence of ^1^O_2_ photosensitizer dyes (MB and RB) and the ^1^O_2_ scavenger His. (A) The effect of HL alone (HL vs. LL), (B) The effect of 5 mM His on HL-induced gene expression, i.e. His +HL vs. HL, abbreviated as (HL+His vs HL). (C) Gene induction by 0.5 µM MB in the presence of HL, i.e. MB+HL vs. HL, abbreviated as (MB vs HL), and (D) the effect of 5 mM His on MB-induced gene expression, i.e. HL+MB+His vs. HL+MB. (E) Gene induction by 0.5 µM RB in the presence of HL, i.e. HL+RB vs. HL, and (F) the effect of 5 mM His on RB-induced gene expression, i.e. HL+RB+His vs. HL+ RB. The statistical significance (-log_10_ p-values) is shown on the Y axis versus the magnitude of change (log_2_fold change) on the X axis. The data points represent the mean value of 3 biological replicates. The expression results of the *hliB* (SGL_RS06470) gene are highlighted in red.

Since HL can induce gene expression via different mechanisms, the ^1^O_2_-specific genes were identified by the addition of 5 mM His which was present during the light treatment. His largely eliminates ^1^O_2_, via chemical scavenging, before it could interact with other molecules. Therefore, the subset of HL-induced genes whose expression is decreased or increased in the presence of His, as compared to the absence of His, represents genes whose expression is specifically enhanced or suppressed by ^1^O_2_, respectively (Fig. 3B, D, F).

Considering that the generation of ^1^O_2_ by photosensitizer dyes requires high light illumination, we used the expression results of the HL treatment as a control for evaluating the gene expression changes induced by using MB or RB to generate exogenous ^1^O_2_ in the cells. In this way the background effect of HL on gene expression could be eliminated. For verification purposes the treatments were carried out both with RB and with MB (separately). In order to avoid unreliable results of the relative expression calculations, genes with negligible read numbers (max group mean < 10) in the respective control samples were disregarded.

Fig. 3C shows those genes whose expression changed in the presence of MB relative to the HL control, whereas Fig. 3D depicts the effect of His addition on the MB-induced effect. Fig. 3E shows those genes whose expression changed in the presence of RB relative to the HL control, whereas Fig. 3F depicts the effect of His addition on the RB-induced effect. As described above for the His+HL vs. HL treatment, the genes in Fig. 3D (His+MB+HL vs. MB+HL) and 3F (His+RB+HL vs. RB+HL) represent the subset of genes whose expression was specifically affected by externally generated ^1^O_2_ induced by the MB and RB photosensitizer dyes, respectively.

### Identification of genes with ^1^O_2_-specific transcript level changes

In order to identify genes whose expression is modified specifically by ^1^O_2_ we created plots by which the affected genes can be organized into specific groups according to their treatment-specific responses (Fig. 4). The set size in Fig. 4A represents the number of genes, which were affected by the indicated treatment either alone or in combination with other treatments (e.g. 80 genes were affected by RB in one way or the other). The intersection size shows the number of genes, which were affected by a particular treatment or by the combination of two or more treatments (e.g. 30 genes were induced by RB alone and additional 25 genes by both RB and MB, etc.). Fig. 4B shows the names of genes, which belong to the most important ^1^O_2_-responsive groups. The 25 genes which were induced by both MB and RB include a transcriptional regulator, ABC transporters, iron homeostasis-related genes, as well genes with unknown function. Genes which were induced by both HL and RB include *psbA2*, providing the bulk of *psbA* transcript for the synthesis of the D1 subunit of the PSII reaction center. The *slr0228* gene, which encodes one member of the FtsH protease family and plays an important role in the degradation of photodamaged D1 protein molecules (Silva *et al*., 2003) was also induced by both HL and RB. The addition of His reversed the HL- and/or MB-, RB -induced upregulation of several genes, which include response regulators, the *isiA* Chl binding protein, and the *sigD* transcription factor (Fig. 4B). Three genes belong to the group whose members can be equally induced by HL, MB, and RB and at the same time the induction by RB can be reversed by His, providing strong support for ^1^O_2_ specificity. We could identify only one gene (*ssr2595*) encoding for the high light inducible *hliB*/*scpD* protein whose transcription was enhanced by HL, MB or RB and at the same time was reversible by His, clearly demonstrating its specificity for ^1^O_2_.

**Figure 4.**
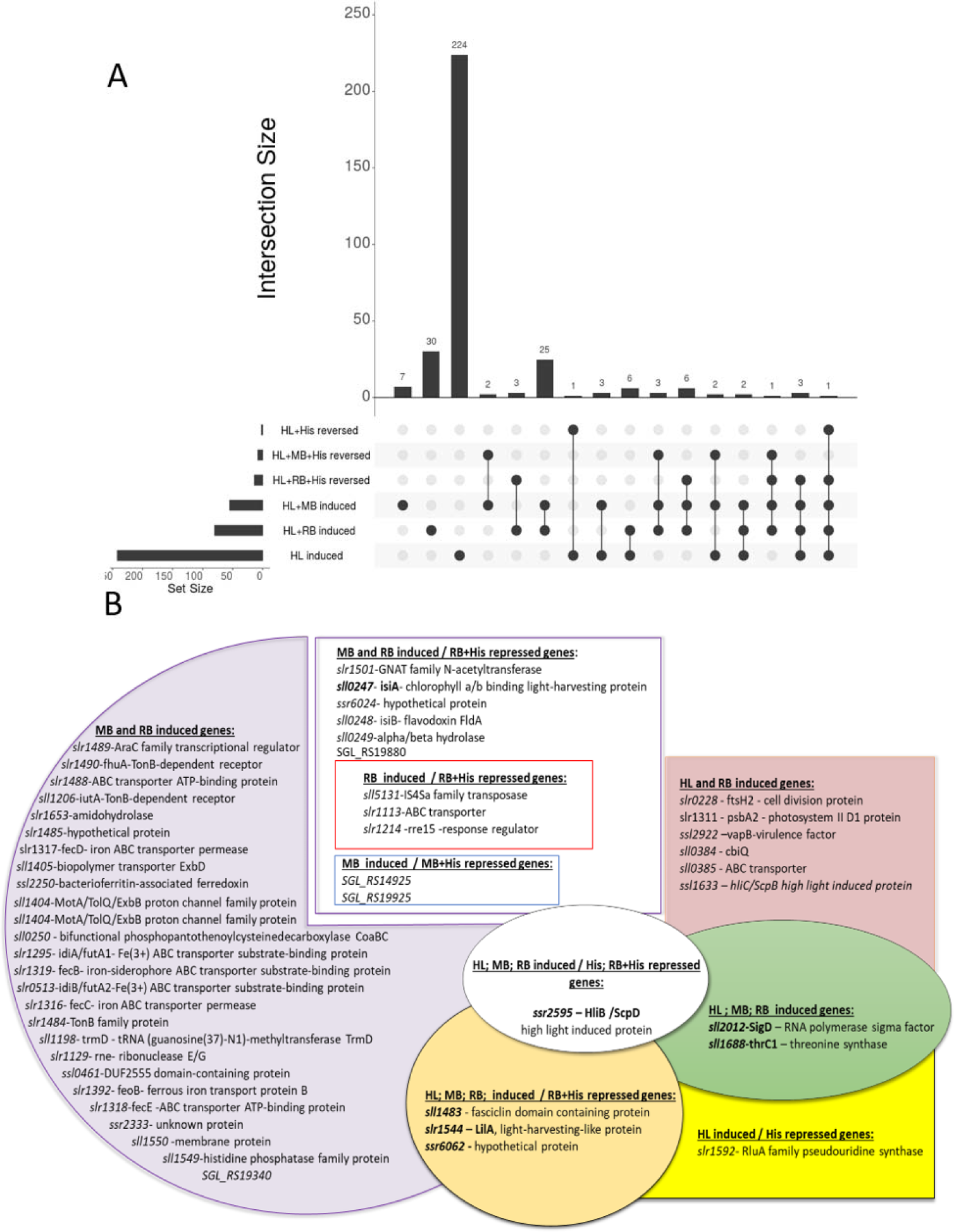
Diagrams of the ^1^O_2_-induced gene sets **A**, Representation of the sizes of ^1^O_2_-induced gene sets, indicating the connection of individual sets and the number of genes belonging to them. Horizontally, the Set Size bars indicate the total number of genes induced/repressed during the given treatment. Vertically, the Intersection Size bars represent the number of genes which belong to the intersections of the various treatments (marked with full dots connected by a line) and the number of genes remaining in the set without intersection by other treatments (indicated by one full dot). **B**, Groups of the induced genes belonging to the most important categories of treatments used for ^1^O_2_ generation. The genes studied in detail are highlighted in bold.

As shown above in Fig. 3, ^1^O_2_ can not only enhance, but also suppress the transcript abundance of several genes. The details of this effect are summarized in Fig. 5. Three genes were suppressed by both RB and HL and three additional genes by both RB and MB. These include the *coaT* and *cadA* P-type ATP-ases, transfer RNAs for different amino acids, a phycobilisome core-rod linker peptide and hypothetical proteins.

**Figure 5.**
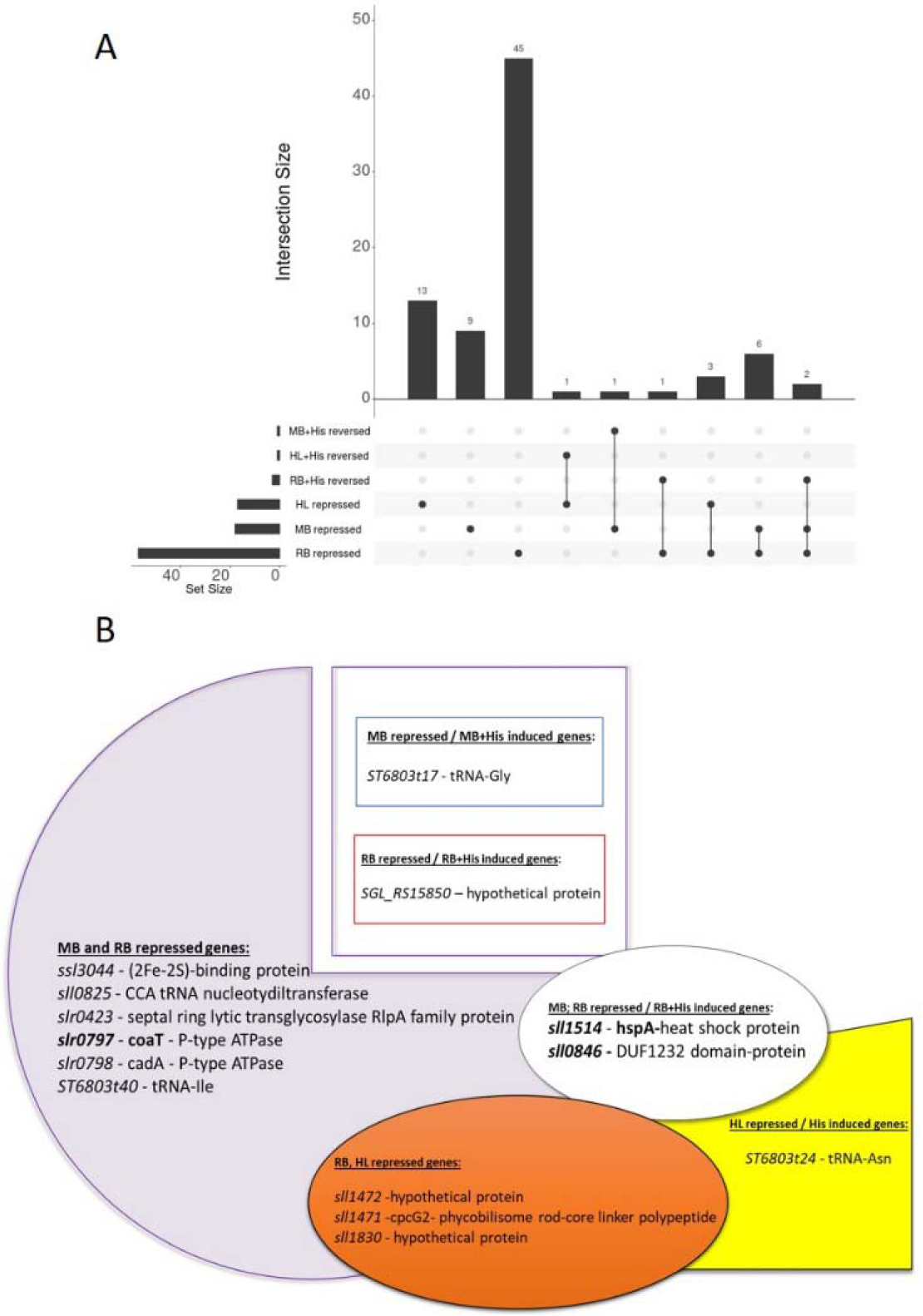
Diagrams of the ^1^O_2_-repressed gene sets **A**, Representation of the sizes of ^1^O_2_-repressed gene sets, indicating the connection of individual sets and the number of genes belonging to them. Horizontally, the Set Size bars indicate the total number of genes whose transcript level changed above the threshold level during the given treatment. Vertically, the Intersection Size bar represents the number of genes which belong to the intersections of the various treatments (marked with full dots connected by a line) and the number of genes remaining in the set without intersection by other treatments (indicated by one full dot). **B**, Groups of the repressed genes belonging to the most important categories of treatments used for ^1^O_2_ generation. The genes studied in detail are highlighted in bold.

^1^O_2_ specificity could be demonstrated by His-induced reversal of the transcript suppression effect in case of several genes, which encode transfer RNAs, the *HspA* heat-shock protein, and a DUF1232 domain protein.

### qPCR confirmation of ^1^O_2_-induced gene expression changes

From the large set of ^1^O_2_-inducible and repressible genes 13 were selected for confirmation by qPCR analysis: the high-light-inducible gene family (*hliA*, *hliB, hliC* and *hliD*), *slr1544* (*lilA*, a light-harvesting-like protein which forms a transcription unit with *hliB*), *sll1483* (a protein with unknown function containing a periplasmic fasciclin domain)*, ssr6062* (hypothetical protein)*, sll1688* (*thrC1*, a threonine synthase), *sigD* (an RNA polymerase sigma factor), *isiA* (iron- and photooxidative stress-responsive Chl binding protein), *hspA* (a small heat shock protein), *sll0846* (encoding a hypothetical DUF domain containing protein), *coaT* (*slr0797* encoding a presumably Co^2+^ exporting P-type ATPase).

The qPCR analysis confirmed the ^1^O_2_-related gene expression changes, which were obtained from the global transcript analysis for *slr1544, sll1483, ssr6062, Thrc1, sigD*, *coaT*, and *isiA,* and showed a partial agreement in the case of *hspA* and *sll0846*. The function and possible relation of these genes to ^1^O_2_ will be discussed in the Discussion section.

### ^1^O_2_ response of the hli gene family

The most promising gene for ^1^O_2_-specific induction was *hliB* marked as *SGL_RS06470* (old tag: *ssr2595*), which was induced not only by HL, as a result of ^1^O_2_ production from endogenous photosensitizers (most likely Chls from the photosynthetic apparatus), but also by illumination in the presence of exogenous photosensitizers RB and MB in a His-reversible way (Fig. 3, 4B, and 7). In addition, *hliB* was one of the genes which showed the strongest increase in expression (log_2_FC = 4.7) upon HL exposure (Fig. 3A). The dependence of *hliB* expression on high light and oxidative stress has already been described (He *et al*., 2001; Akulinkina *et al*., 2015; Cheregi & Funk, 2015; Konert *et al*., 2022), but the involvement of ^1^O_2_ in the regulation of *hliB* expression has not been known.

Since *hliB* belongs to the *hli* gene family, all of whose members are induced by high light (Bhaya *et al*., 2002; Cheregi & Funk, 2015; Konert *et al*., 2022), we aimed to verify whether or not the other *hli* genes (*hliA*, *hliC* and *hliD*) also respond to ^1^O_2._ To this end, we verified the transcript level changes of the four *hli* genes using qPCR. The data show that the other three members of the *hli* gene family respond similarly to HL, RB, MB as *hliB*, although the extent of their response is smaller than that of *hliB*, especially in case of *hliD* (Fig. 7A). Since the induction of the transcript levels is largely prevented by His, one can conclude that ^1^O_2_ is involved in the regulation of the expression of all *hli* genes, although the strongest response is shown by *hliB*. As was shown in Fig. 6 above, *lilA* (*slr1544*), which forms a common transcription unit with *hliB* also shows a strong upregulation by HL, RB, and MB in a His-reversible way, therefore it is also identified as a ^1^O_2_-responsive gene.

**Figure 6.**
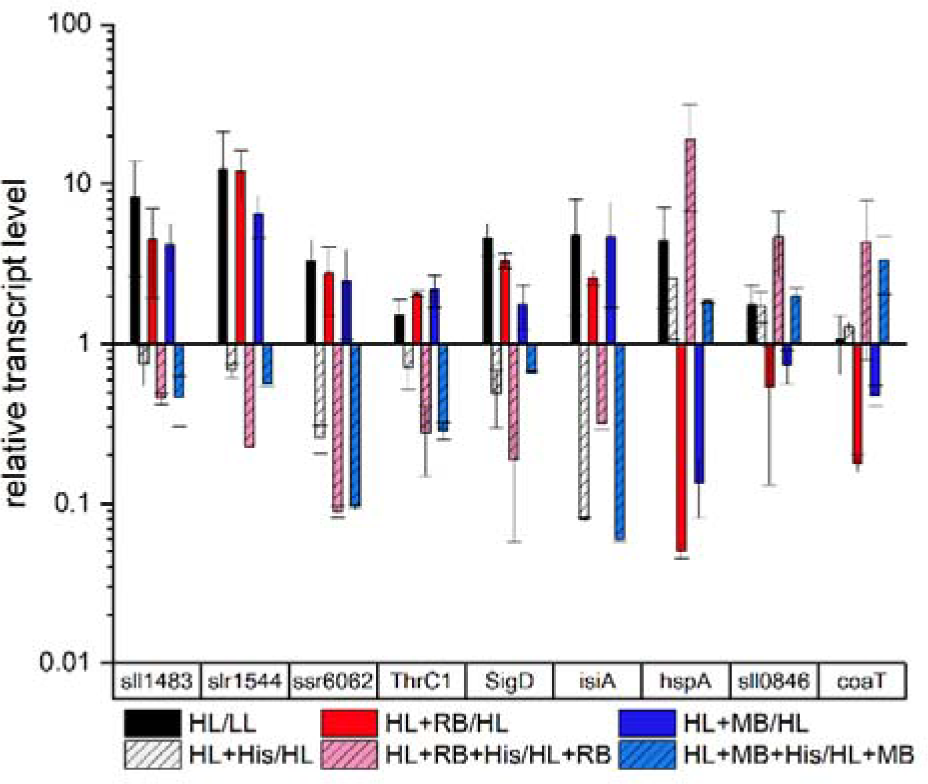
Gene expression of genes selected for detailed investigation by qPCR. Mean+SE, n=3).

**Figure 7.**
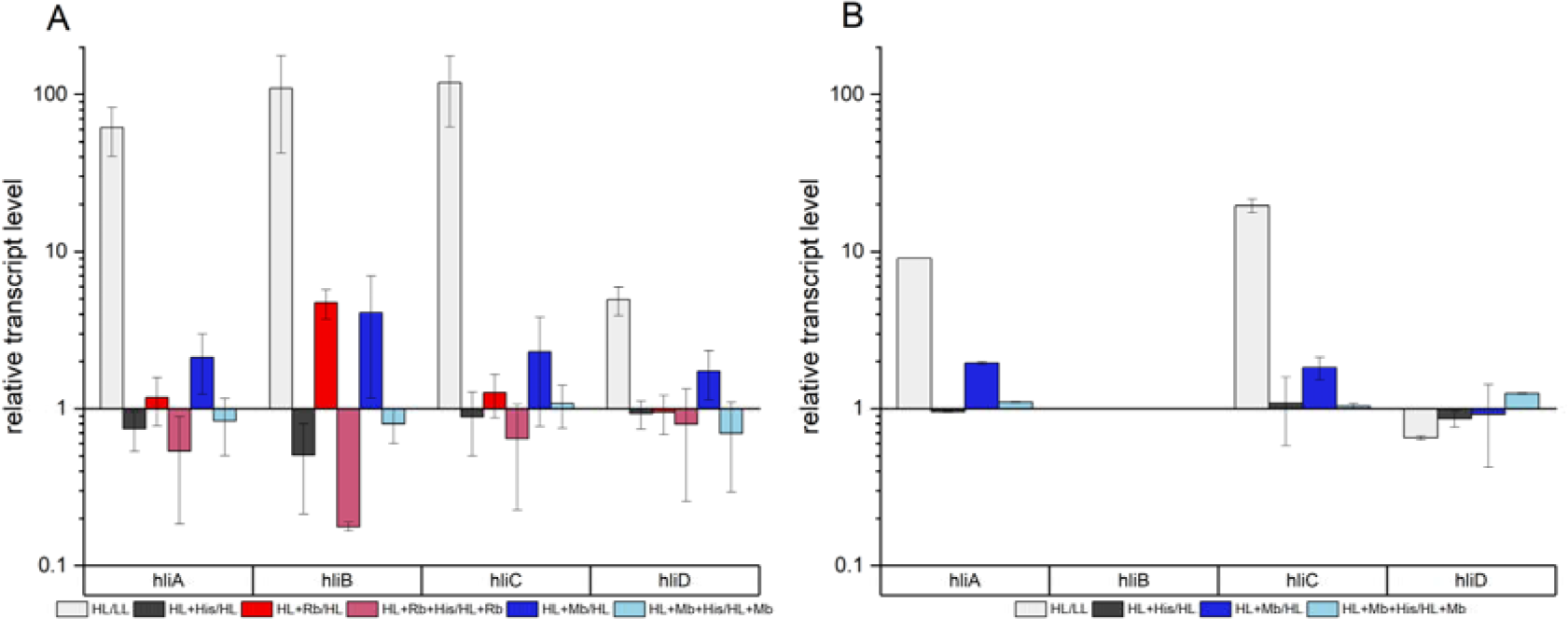
Expression changes of *hli* genes in WT (A) or Δ*hliB* (B) *Synechocystis* cells exposed to LL and HL illumination with 0.5 µM MB and its application with 5 mM His (Mean+SE, n=3).

In order to clarify if the *hliB* gene or its HliB protein product might be involved in the regulation of the other members of the *hli* gene family, we created a Δ*hliB* mutant via replacing this gene by a *Spectinomycin* resistance cassette (Fig. 1). In this deletion mutant the ^1^O_2_-induced expression of the *hliA* and *hliC* genes was practically the same as in the WT. Interestingly, however, *hliD* was not expressed in the deletion mutant under HL or MB exposure (Fig. 7B). These data show that *hliB* deficiency has no influence on the transcript levels of *hliA* and *hliC*, but has a direct or indirect influence on the expression of *hliD*.

### The protective role of HliB against ^1^O_2_-mediated photodamage

The ^1^O_2_-specific enhancement of *hliB* gene expression raises the possibility that the HliB protein has a well-defined role in protection against ^1^O_2_-induced damage. In order to clarify this hypothesis, the WT and Δ*hliB* mutant strains were grown under HL illumination in the absence and presence of the ^1^O_2_ sensitizers MB or RB. Both dyes resulted in a significant growth inhibition in the WT strain, which was more pronounced in the presence of MB then RB, demonstrating the overall damaging effect of exogenous ^1^O_2_ on cell growth. Deletion of the *hliB* gene enhanced the growth retardation not only in the presence of both dyes, but also under HL without addition (Fig. 8). This phenotype of the Δ*hliB* mutant supports the assumption that HliB plays an important role in the defense against ^1^O_2_ in *Synechocystis*. These data are in basic agreement with earlier results showing HL-induced growth retardation of a *Synechocystis* mutant lacking all the four *hli* genes (He *et al*., 2001; Havaux *et al*., 2003). However, our data also demonstrate that the deletion of *hliB* alone is sufficient for growth retardation, and show that ^1^O_2_ plays an important role in HL-induced growth retardation in the absence of HliB.

**Figure 8.**
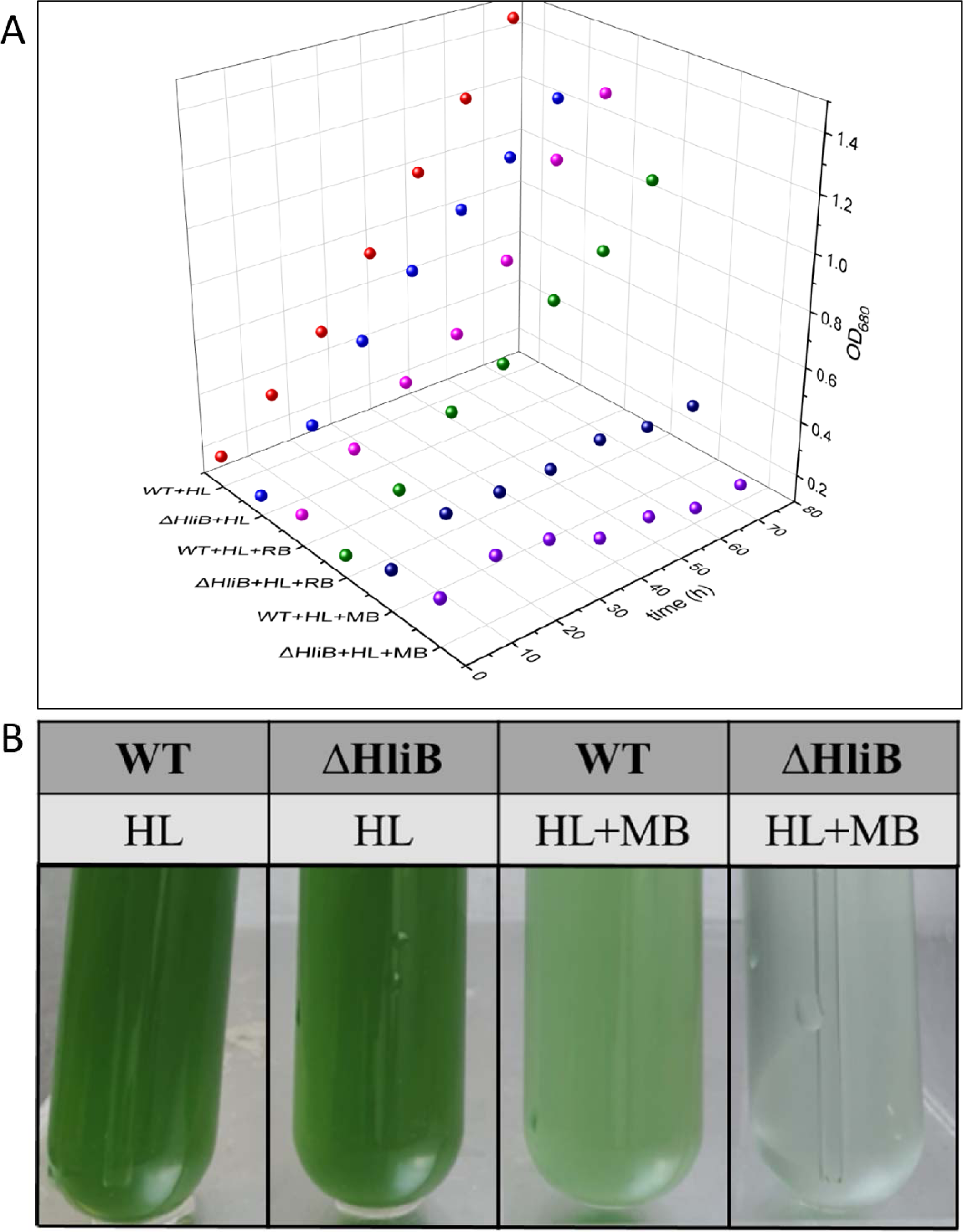
**A:** The growth curves of the WT and Δ*hliB Synechocystis* cells in different ^1^O_2_ treatments. **B:** The photo of visible growth inhibition of Δ*hliB* and WT *Synechocystis* cells under HL illumination in the continuous presence of MB (0.5 µM).

### The hliBLux ^1^O_2_ bioreporter strain

An important reason for carrying out the global transcriptome survey was to find promoters that respond specifically to ^1^O_2_, allowing the construction of whole cell bioreporters, similarly to our previous studies (Peca *et al*., 2007, 2017; Patyi *et al*., 2021). Such a strain would facilitate continuous monitoring of ^1^O_2_ production, a technique that was so far not available for cyanobacteria. The results described above show that the *hliB* gene is specifically induced by ^1^O_2_. In order to utilize its ^1^O_2_-dependent expression we constructed a *Synechocystis* strain by fusing the *hliB* promoter with the bacterial luciferase reporter system, and designated it as *hliB*Lux. To test its utility for ^1^O_2_ sensing in cellular environment we performed ^1^O_2_ treatments using the same approach as for the gene expression studies, but instead of transcript levels the bioluminescence intensity was detected (Fig. 9).

**Figure 9.**
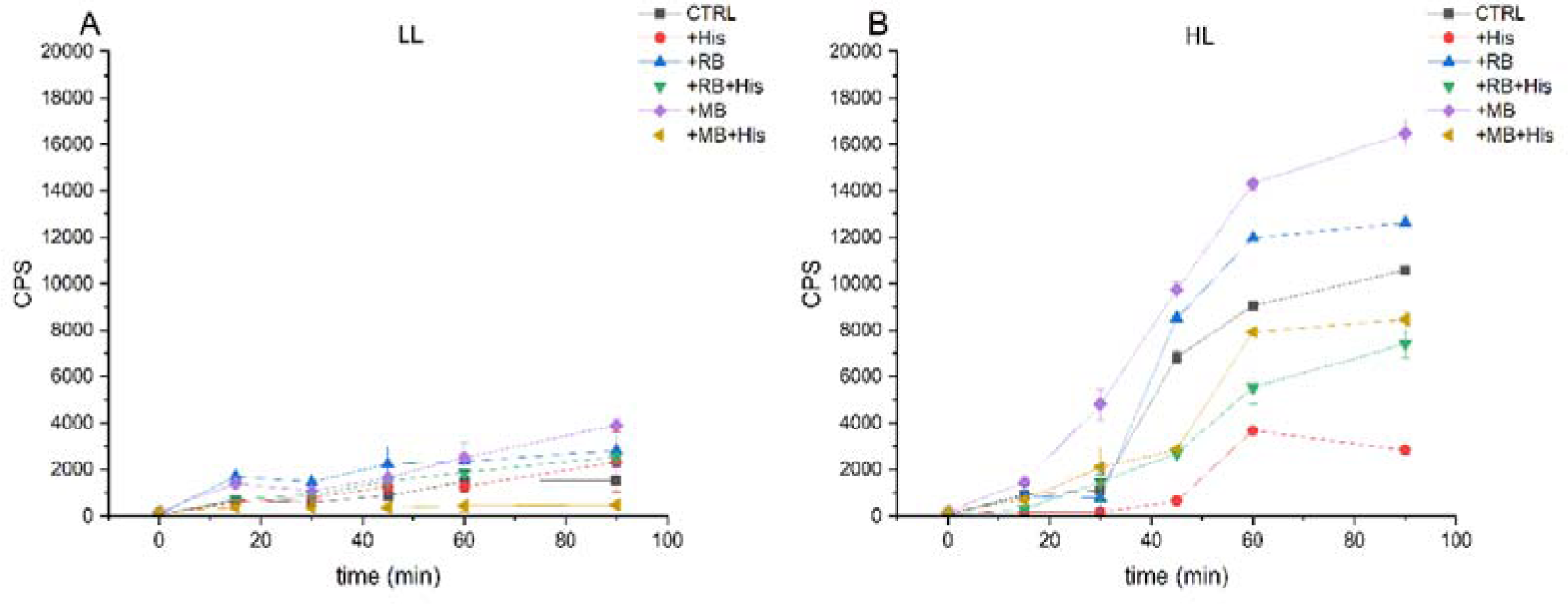
The bioluminescent response of the *hli*Blux strain to MB and RB. Cells were incubated in BG-11 medium exposed to LL (A) or HL (B) conditions and supplemented with different photosensitizers with or without 5 mM His before bioluminescence was measured. Each point represents the mean of three parallels.

When *hliB*Lux cells were exposed to LL, only a moderate luminescence response was observed (Fig. 9A). However, the luminescence response was enhanced by HL ca. 7-fold relative to the LL conditions (Fig. 9B). Importantly, the luminescence induction was largely retarded in the presence of His showing that it was caused by endogenously produced ^1^O_2_ (Fig. 9B). The luminescence intensity was further increased when HL exposure was performed in the presence of MB or RB, and His addition retarded both effects (Fig. 9B). These data provide evidence that the luminescence response indeed reflects ^1^O_2_ production and also that the *hliB*Lux strain can be used to detect ^1^O_2_ inside the *Synechosystis* cells when produced from either endogenous or exogenous sources.

Even though MB and RB were applied at the same (0.5 μM) concentration, MB produced higher luminescence response of the *hliB*Lux biosensor than RB (Fig. 9) in agreement with the higher level *hilB* upregulation by MB (Figs. 3C, 3E). On the other hand, RB induced a higher number of genes than MB (Figs. 3C, 3E and 4A) both in the group where His was able to reverse the induction effect and in the groups where the His effect was not obvious. This might be related to the different accessibility of the interior of *Synechocystis* cells for the two dyes and/or to the accessibility for His of the cell compartments that MB or RB could reach and where they could produce ^1^O_2_.

## Discussion

In the present work we performed a complete transcript analysis in the cyanobacterium *Synechocystis* 6803 under conditions where ^1^O_2_ was generated either from endogenous (HL-excited natural pigments) or exogenous (HL-excited MB or RB photosensitizers) sources. Our transcript profile analysis revealed a large set of ^1^O_2_–responsive genes, which demonstrates that cyanobacteria are able to respond to ^1^O_2_ exposure by specific changes in gene expression, similarly to all other photosynthetic organisms.

### Overlap of ^1^O_2_- and HL-dependent regulation of gene induction in Synechocystis

In photosynthetic organisms ^1^O_2_ is produced mostly via the Type-II photodynamic effect of pigments which participate in harvesting light energy. In addition, ^1^O_2_ production via the photodynamic effect is practically linearly proportional with light intensity (Rehman et al., 2013). Therefore, one would expect to find ^1^O_2_-induced upregulation of genes mostly among the high-light-inducible genes of *Synechocystis*. Previous studies have identified a large set of genes with light-dependent expression profile in *Synechocystis* (He *et al*., 2001; Hihara *et al*., 2001; Bhaya *et al*., 2002; Hsiao *et al*., 2004; Ogawa *et al*., 2018). However, the possible involvement of ^1^O_2_ in this effect has not been studied previously. Our analysis has indicated the presence of 82 HL-responsive genes whose expression change is prevented by the ^1^O_2_ scavenger His (Fig. 3B) and therefore can be assigned to ^1^O_2_.

Intracellular ^1^O_2_ production by photosensitizer dyes such as RB or MB also induced several genes in a His-reversible way, which showed a partial overlap with the HL-inducible ^1^O_2_-dependent genes (Figs. 3D, 3F, 4). From these data we can safely conclude that HL-induced regulation of a sizeable subset of *Synechocystis* genes is mediated, at least partly, by ^1^O_2_-dependent signaling pathways.

### ^1^O_2_-mediated regulation of the hli gene family

The *hli* family is of a special importance among the ^1^O_2_-responsive genes. It consists of 4 members (A-D) which encode small chlorophyll-binding proteins, also called SCPs, or HLIPs. There is a 5^th^ gene (*slr1544* or *lilA*) which is structurally similar to the SCPs and forms a transcription unit with *hliB* (Kufryk *et al*., 2008). Both the SCP and LilA proteins contain a long hydrophilic N-terminal region and a transmembrane C-terminal region, with two residual conserved regions of Chl-binding domains. It has also been shown that LilA can bind to PSII under stress (Kufryk *et al*., 2008) similarly to HilB and HliC (Komenda & Sobotka, 2016). Our data demonstrate that ^1^O_2_ is involved in the HL-induced upregulation of the *hli* and *lilA* genes (Figs. 6 and 7). It is interesting to note that while the *hli* genes are expressed only under HL exposure (and some other stress conditions such as low temperature, N- and S-limitation (He *et al*., 2001)) in WT *Synechocystis* cells, they are constitutively present in the PSI-less mutant (Funk & Vermaas, 1999). This observation can be explained by the increased rate of ^1^O_2_ formation in the PSI-less mutant (Kodru *et al*., 2020), which induces a high level of *hli* transcripts.

^1^O_2_ production and the related photodamage is enhanced in the *Synechocystis* mutants lacking the *hli* and *lilA* genes demonstrating the important protective role against ^1^O_2_-mediated photodamage of their protein products (He *et al*., 2001; Sinha *et al*., 2012). Our data show that the lack of the most strongly ^1^O_2_-responsive *hli* gene, *hliB*, in itself is sufficient to induce significant retardation of cell growth under conditions of ^1^O_2_ production (Fig. 8). In agreement with previous suggestions (Sinha *et al*., 2012) the protective role of the Hli proteins can be exerted by binding Chl molecules, which are released during photodamage and/or repair of the PSII complex and represent a dangerous source of ^1^O_2_ production in their free form, in a protein environment which is similar to that of the light-harvesting complexes and includes protective carotenoids (Konert *et al*., 2022) that decrease ^1^O_2_ formation. It is of note that the HliB protein was also found in association with PSII (Promnares *et al*., 2006; Komenda & Sobotka, 2016).

Although all the four *hli* genes were induced by ^1^O_2_, the extent of their upregulation was different, *hliB* (and the co-transcribed *lilA*) and *hliD* being the most and the least induced ones, respectively. The reason for this difference is not fully clear, but most likely related to the promoter region of the genes which may have specific sites for sensing a transcription factor or some other component that is produced or modified by ^1^O_2_ (Page *et al*., 2017).

Deletion of the *hliB* gene had only a minor effect on the ^1^O_2_ response of *hliA* and *hliC*, but largely eliminated the induction of *hliD* (Fig. 7B). These data show that the HliB protein does not participate in a signaling event that would influence the ^1^O_2_-dependent expression of *hliA* and *hliC*; however, it has an indirect or direct effect on the expression of *hliD*.

We have utilized the strong, ^1^O_2_-specific induction of *hliB* to create a ^1^O_2_ reporter construct that contains the ^1^O_2_-inducible *hliB* promoter fused to bacterial luciferase. It responds to ^1^O_2_ when generated either by the endogenous pigments of the photosynthetic apparatus or by added photosensitizer dyes in a His-repressible manner (Fig. 9), and provides a useful tool for monitoring intracellular ^1^O_2_ conditions under various stress conditions.

### Effect of ^1^O_2_ on Photosystem II repair-related genes

Repair of photodamaged PSII is a very important process that helps to maintain photosynthetic activity under elevated light conditions (Aro *et al*., 1993). The main steps of PSII repair are proteolytic degradation of the damaged D1 subunits via involvement of the FtsH protease (Silva *et al*., 2003), followed by *de novo* synthesis of new D1 copies which is regulated transcriptionally in cyanobacteria by producing *psbA* mRNA (Nixon *et al*., 2010; Järvi *et al*., 2015). In order to function optimally, the repair process should be closely regulated not only by the already occurred damage, but also by the factors that induce the damage to PSII. Therefore, it is not surprising that ^1^O_2_, which is an important damaging factor of PSII structure and function (Vass & Cser, 2009; Zavafer, 2021) induces upregulation of genes which are involved in PSII repair, such as *psbA2* (Fig. 4, Supplementary Table 1) that provides the largest contribution to the *psbA* transcript pool for synthesis of D1 (Mohamed *et al*., 1993). Interestingly, both the *ftsH2* and *ftsH3* genes, which encode the FtsH2 and FtsH3 subunits of the D1-degrading FtsH complex are upregulated by ^1^O_2_. These data show that the well-known HL-induced upregulation of the *ftsH2* and *ftsH3* and *psbA2* genes (see e.g. Hernández-Prieto et al., 2016) occurs partly via ^1^O_2_, which helps to induce the PSII repair machinery on the side of both D1 degradation and de novo synthesis.

The ^1^O_2_-induced boosting of PSII repair is particularly interesting, since a long-standing dogma states that ^1^O_2_ does not induce damage to PSII, therefore would not need to enhance its repair, and the only effect of ^1^O_2_ in PSII photoinhibition would be the inhibition of *de novo* D1 synthesis (Nishiyama *et al*., 2004), which contradicts various observations (Rehman *et al*., 2013; Fischer *et al*., 2013; Bashir *et al*., 2021) including our present findings.

### Overlap of ^1^O_2_ with the salt-, hyperosmotic and cold stress response

Salt-, hyperosmotic and cold stress are characteristic phenomena, which change the expression of a large set of genes (Kanesaki *et al*., 2002; Liu *et al*., 2013) and can enhance photodamage. Therefore, it is interesting to note that some of the ^1^O_2_-responsive genes identified in our work are also induced by salt and/or hyperosmotic or cold-stress conditions. The most important genes, which are upregulated by ^1^O_2_ as well as by salt- and/or hyperosmotic stress are *hliA*, *hliB*, *lilA*, *isiA, ssr2016*, *sll1483* (fasciclin), and *sll1515* (DUF4278) based on the comparison of our data and earlier published results (Kanesaki *et al*., 2002). Some other genes (e.g. *hliA*, *hliB*, *hliC*, *psbA2*, and tRNAs for several amino acids) are shared between the ^1^O_2_- and cold-induced transcript changes based on the comparison of our data with earlier published results (Liu *et al*., 2013).

These observations point to the possibility that ^1^O_2_ may participate in the regulation of the expression of the above genes not only under HL, but also under other stress conditions (salt, hyperosmotic, cold), or that these genes can be regulated independently by several mechanisms. Clarification of this interesting point will require further studies.

### Overlap of ^1^O_2_ with iron homeostasis

In *Synechocystis* and some other cyanobacteria, iron starvation leads to the synthesis of the IsiA protein which protects the photosystems against oxidative stress. *isiA* encodes a Chl-containing light harvesting complex and has been shown to be upregulated not only by iron starvation but also by high light under iron replete conditions (Havaux *et al*., 2005). We have observed His-reversible induction of *isiA* by HL and RB or MB (Fig. 6), which points to the involvement of ^1^O_2_ in the HL-induced response of *isiA*.

One of the suggested functions of *isiA* is to provide temporary binding environment for Chls which are released from light harvesting complexes under Fe-limiting conditions (Riethman *et al*., 1988; Chen *et al*., 2018). Our observation of ^1^O_2_-induced upregulation of *isiA* confirms this idea, since free Chls are very efficient ^1^O_2_ sensitizers whose presence should be avoided. Therefore, IsiA could exert photoprotection via the same mechanism as the HLIP (SCP) proteins, i.e. by providing a temporary sequestration for Chls in the vicinity of protective carotenoids. It has also been suggested that IsiA forms a large photoprotective complex together with HliA and some other proteins, as well as carotenoids (zeaxanthin and myxoxanthophyll) (Daddy *et al*., 2015).

Oxidative stress in general has been implicated in the induction of *isiA* since it was induced in *Synechocystis* under iron replete conditions not only by high light, but also by salt stress, methylviologen and H_2_O_2_ (see (Michel & Pistorius, 2004) and (Havaux *et al*., 2005) and references therein). Our data show that one of the specific oxidative signals that induce *isiA* expression is ^1^O_2_, but this does not exclude the involvement of other oxidative components which may act independently of, or indirectly via, ^1^O_2_. Besides *isiA* the iron-deficiency-induced *idiA* gene was also induced by RB treatment, which is in agreement with the earlier suggested role of IdiA in protection against photooxidative damage (Glaeser *et al*., 2011).

### ^1^O_2_ induced suppression of Synechocystis genes

A potentially interesting finding of our work was the identification of genes, which were suppressed by ^1^O_2_. This aspect of ^1^O_2_-induced gene responses had been realized earlier in *Arabidopsis,* where ^1^O_2_ induced the suppression of genes involved in the synthesis of photosynthetic pigments (Page *et al*., 2017), which was, however, not observed in our case. In *Synechocystis* the *sll1471*, *sll1472* and *sll1830* genes were suppressed by HL and RB in a partly His-reversible way, indicating the involvement of ^1^O_2_. *sll1472* and *sll1830* encode unknown proteins; however, the gene product of *sll1471* is the so-called CpcG2, a linker protein for the Photosystem I phycobilisome antenna. CpcG2 has been shown to participate in the formation of the NDH-1L-CpcG2-PSI supercomplex that facilitates cyclic electron transport around PSI via NDH-1L in cyanobacteria (Gao *et al*., 2016), whose absence leads to high light sensitivity. Since cyclic electron flow is a protective mechanism against HL-induced stress, which also protects against ^1^O_2_ formation, it would be physiologically counterintuitive to assume ^1^O_2_-induced downregulation of CpcG2 leading to decreased cyclic electron flow as part of photoprotection. Therefore, it is more probable that the suppression of *sll1471* (CpcG2) expression is due to a damaging effect of ^1^O_2_.

Some other genes such as *coaT* (*slr0797*), *cadA* (s*lr0798*), *ssl3044*, *sll0825*, *sll0423*, *tRNA-Ile*, *sll1514*, s*ll0846*, and *RS1585*0 were also suppressed by exogenously generated ^1^O_2_ in a His-reversible way, but at the same time were induced by HL, showing a tendency for His reversibility. This apparently contradicting behaviour can probably be explained by a damaging effect of externally generated ^1^O_2_, which has a higher rate than that induced by HL alone and reverses the HL-induced upregulation of these genes. It is, however, unclear why only few of the ^1^O_2_-responsive genes show this effect.

### ^1^O_2_-related signal transduction pathways

Due to the short lifetime of ^1^O_2_, it is very likely that the sensor for ^1^O_2_ is located in close proximity to the source of ^1^O_2_ in the thylakoids. Possible candidates in *Chlamydomonas* could be the degradation products of the D1 protein, or lipid peroxides (Fischer *et al*., 2006). As regards the downstream elements of signal transduction, a small zinc finger protein METHYLENE BLUE SENSITIVITY (MBS) has been identified that is required for induction of ^1^O_2_-dependent gene expression in *Chlamydomonas* (Shao *et al*., 2013). In *Chlamydomonas* a strong ^1^O_2_-specific upregulation was also observed for the cytosolic glutathione peroxidase homolog gene (*GPXH*/*GPX5*) (Leisinger *et al*., 2001; Fischer *et al*., 2006; Ledford *et al*., 2007). This effect was abolished by the deletion of the *psbP2* gene, which encodes a small protein that is similar to the PsbP subunit of the water-oxidizing complex of PSII, and may act as a mediator of the signal from ^1^O_2_ generated in the chloroplast (Brzezowski *et al*., 2012). It has also been shown that Chl precursors such as Mg-protoporphyrin IX can act as signaling molecules between chloroplasts and the nucleus (Strand *et al*., 2003). By analogy, it can be assumed that Chl degradation products such as pheophytin, chlorophyllide or pheophorbide can act as signaling molecule (Krieger-Liszkay, 2005).

In higher plants (*Arabidopsis*) β-carotene, residing in the PSII core complex, has also been suggested to act as a ^1^O_2_ sensor. In this model the ^1^O_2_-specific endo-peroxides, such as β-cyclocitral (Ramel *et al*., 2012), which are highly reactive, volatile and electrophilic compounds produced via ^1^O_2_-mediated oxidative modification of β-carotene can induce gene expression changes (Dogra & Kim, 2020). It has also been shown that the *Chlamydomonas METHYLENE BLUE SENSITIVITY* (*MBS*) gene has homologs in *Arabidopsis* (Shao *et al*., 2013), and one of them (*MBS1*) acts in ^1^O_2_ signaling downstream of β-cyclocitral (Shumbe *et al*., 2017).

In *Arabidopsis* the so called EXECUTER1 and EXECUTER2 proteins are also implied as important mediators of ^1^O_2_-induced gene expression (Keun *et al*., 2007). According to more recent findings the Executer proteins may undergo structural modifications as a result of ^1^O_2_ attack, which makes possible a proteolytic degradation and the production of smaller fragments that can serve as signaling components (Dogra & Kim, 2020). It has also been shown that the ^1^O_2_-modified EXCUTER1 is degraded by the FtsH2 protease (Wang *et al*., 2016), which also plays a key role during PSII repair by degrading the D1 subunit of the PSII (Bailey *et al*., 2002) whose photodamage also involves ^1^O_2_ (Mizusawa *et al*., 2003; Vass, 2012).

The best explored situation is in *Rhodobacter sphaeroides* and other bacteria, such as *E. coli*, where ^1^O_2_-induced gene expression is controlled by the alternative group IV sigma factor σE (or RpoE) and its cognate anti-sigma ChrR. Under normal conditions RpoE and ChrR form a complex, which prevents RpoE from interacting with RNA polymerase. After interaction with ^1^O_2_ ChrR is degraded and releases RpoE that can induce gene expression (Anthony *et al*., 2005; Campbell *et al*., 2007; Nuss *et al*., 2013). *Synechocystis* has two rpoE homologues, *sigH* (sll0856) and *sigG* (slr1545) (Huckauf *et al*., 2000). However, the expression of these genes was not affected by ^1^O_2_ in our experiments in contrast to the response of *Rhodobacter rpoE*, which was strongly induced by ^1^O_2_ (Anthony *et al*., 2005). So far we could not find an obvious homolog of *chrR* in *Synechocystis*, therefore there is no obvious evidence for the presence of an RpoE-dependent ^1^O_2_ signaling pathway in *Synechocystis*.

It is of note, however, that among the 9 sigma factors of *Synechocystis* there was one, the Group-2 *sigD* (*sll2012*), which was induced by HL as well as RB and MB (Fig. 4 and 6). The SigD protein is involved in several stress responses, for example in regulation related to the Hik response regulators (Shoumskaya *et al*., 2005; Los *et al*., 2010). Induction of the *sigD* gene was partially suppressed by His in the case of HL, RB and MB (Supplementary Table 1). This finding strongly indicates that ^1^O_2_ is involved in the expression of *sigD*, which appears to be the only ^1^O_2_-dependent sigma factor in *Synechocystis*. *sigD* expression has been shown to be upregulated by light (Imamura *et al*., 2003), as well as by HL and heat stress (Turunen *et al*., 2022), both of which can involve ^1^O_2_ formation (for heat-induced ^1^O_2_ formation see (Prasad *et al*., 2016)), providing support for our finding. However, the deletion of *sigD* in *Synechocystis* did not eliminate the HL-induced upregulation of *hliB*, *lilA,* or *ssr2016* (coding for a hypothetical protein) (Turunen *et al*., 2022), which are among the most strongly ^1^O_2_-responsive genes according to our results. HL-induced *hliB*, *lilA,* or *ssr2016* expression was not affected by deleting *sigB*, *sigBCE* or *sigCDE* either (Turunen *et al*., 2022), which shows that Group-2 sigma factors are unlikely to participate in ^1^O_2_-dependent signaling in *Synechocystis*.

It is important to note that all *hli*/*scp* genes have the so-called High Light Regulatory 1 (HLR1) element (Eriksson *et al*., 2000) in their promoter regions (Kappell *et al*., 2006), which is also present in some other high-light-inducible genes. Out of the 19 genes with known HLR1 elements (Eriksson *et al*., 2000)(Cheregi & Funk, 2015) 17 were found to be ^1^O_2_-responsive according to our data (*hliA-D*, *psbA2*, *nblA*, *sigD*, and the *sll1483*, *sll0157*, *slr0320*, *slr1687*, *ssl3044* and *ssl2162* genes encoding hypothetical proteins). Only *psbA3* and *slr1894* are absent from our data, which may be due to their low abundance under our conditions. It has also been shown that the factor that binds to the HLR1 motif upstream of the *hliB* gene is RpaB (slr0947), a response regulator (Kappell & Van Waasbergen, 2007), which is expected to modulate the function of the other genes containing HLR1 as well. The current model for the regulation through RpaB, in genes such as the *hli* family where HLR1 is located upstream of the core promoter region, is that under normal and low light conditions binding of the phosphorylated form of RpaB (RpaB-P) to HRL1 prevents the interaction of the core promoter with RNA polymerase. At high light intensities the sensory histidine kinase Hik33 mediates dephosphorylation of RpaB-P, which facilitates the release of RpaB from the HLR1 motif leading to the de-repression of the genes (Hanaoka & Tanaka, 2008; Cheregi & Funk, 2015). Our data did not show any significant change in the abundance of *rpaB* mRNA in response to ^1^O_2_. Therefore, one possibility for the ^1^O_2_-induced upregulation of the *hli* and other genes having the HLR1 motif is that ^1^O_2_-induced damage of the RpaB-P protein induces its release from HLR1, perhaps via a proteolytic degradation step as occurs in case of the ChrR anti-sigma factor in *Rhodobacter sphaeroides* (Nuss *et al*., 2013). This effect could be independent of Hik33, since we did not observe a significant expression change of the gene encoding Hik33; however, other regulation pathways cannot be excluded either at the present stage of knowledge. Since the HLR1 motive is present only in a fraction of the ^1^O_2_-inducible genes, the hypothesized RpaB-dependent regulatory mechanism cannot be a general way of regulating ^1^O_2_-dependent gene expression, therefore extensive future studies will be needed to clarify the molecular basis of how ^1^O_2_ can regulate gene expression in *Synechocystis* and other cyanobacteria.

## Abbreviations

Chl: Chlorophyll
BChl: bacteriochlorophyll
HL: high light
HLIP: high-light-inducible protein
LL: low light
MB: Methylene blue
^1^O_2_: Singlet oxygen
PSII: Photosystem II
qPCR: quantitative PCR
RB: Rose Bengal
SCP: Small chlorophyll binding protein

## Author contributions

IV conceived the idea and conceptualized the study in collaboration with PBK. GP, PBK, IM and BH set up and performed the transcript profiling and RT PCR experiments. PBK, GM and GP analyzed the transcript changes. IM created the deletion mutants, GP created the bioreporter strain and performed the bioluminescence experiments. GP prepared the figures and wrote the first draft of the paper with in-depth suggestions from PBK. IV finalized the conclusions and the text of the paper. All the authors approved the final version of the manuscript.

## Data availability

The data that support the findings of this study are openly available in Supporting Information.

## Supporting Information (brief legends)

S1 file: Selected transcript table

S2 file: Full transcript table

## References

Akulinkina D V., Bolychevtseva Y V., Elanskaya I V., Karapetyan N V., Yurina NP. 2015. Association of high light-inducible HliA/HliB stress proteins with photosystem 1 trimers and monomers of the cyanobacterium Synechocystis PCC 6803. Biochemistry (Moscow) 80: 1254–1261.

Anthony JR, Warczak KL, Donohue TJ. 2005. A transcriptional response to singlet oxygen, a toxic byproduct of photosynthesis. Proceedings of the National Academy of Sciences of the United States of America 102: 6502–6507.

Aro EM, McCaffery S, Anderson JM. 1993. Photoinhibition and D1 protein degradation in peas acclimated to different growth irradiances. Plant Physiology 103: 835–843.

Bailey S, Thompson E, Nixon PJ, Horton P, Mullineaux CW, Robinson C, Mann NH. 2002. A critical role for the Var2 FtsH homologue of Arabidopsis thaliana in the photosystem II repair cycle in vivo. Journal of Biological Chemistry 277: 2006–2011.

Bashir F, Rehman AU, Szabó M, Vass I. 2021. Singlet oxygen damages the function of Photosystem II in isolated thylakoids and in the green alga Chlorella sorokiniana. Photosynthesis Research 149: 93–105.

Berghoff BA, Glaeser J, Nuss AM, Zobawa M, Lottspeich F, Klug G. 2011. Anoxygenic photosynthesis and photooxidative stress: A particular challenge for Roseobacter. Environmental Microbiology 13: 775–791.

Bhaya D, Dufresne A, Vaulot D, Grossman A. 2002. Analysis of the hli gene family in marine and freshwater cyanobacteria. FEMS Microbiology Letters 215: 209–219.

Brzezowski P, Wilson KE, Gray GR. 2012. The PSBP2 protein of Chlamydomonas reinhardtii is required for singlet oxygen-dependent signaling. : 1289–1303.

Campbell EA, Greenwell R, Anthony JR, Wang S, Lim L, Das K, Sofia HJ, Donohue TJ, Darst SA. 2007. A Conserved Structural Module Regulates Transcriptional Responses to Diverse Stress Signals in Bacteria. : 793–805.

Chen J, Shi W, Li W, Chen G, Qin S. 2018. Specific genetic variation in two non-motile substrains of the model cyanobacterium Synechocystis sp. PCC 6803. Journal of Oceanology and Limnology 36: 2322–2332.

Cheregi O, Funk C. 2015. Regulation of the scp Genes in the Cyanobacterium Synechocystis sp. PCC 6803 What is New? Molecules 20: 14621–14637.

Daddy S, Zhan J, Jantaro S, He C, He Q, Wang Q. 2015. A novel high light-inducible carotenoid-binding protein complex in the thylakoid membranes of Synechocystis PCC 6803. Scientific Reports 5: 1–8.

Dmitrieva VA, Tyutereva E V., Voitsekhovskaja O V. 2020. Singlet oxygen in plants: Generation, detection, and signaling roles. International Journal of Molecular Sciences 21.

Dogra V, Kim C. 2020. Singlet Oxygen Metabolism: From Genesis to Signaling. Frontiers in Plant Science 10: 1–9.

Eriksson J, Salih GF, Ghebramedhin H, Jansson C. 2000. Deletion mutagenesis of the 5’ psbA2 region in Synechocystis 6803: Identification of a putative cis element involved in photoregulation. Molecular Cell Biology Research Communications 3: 292–298.

Ferretti U, Ciura J, Ksas B, Rác M, Sedlářová M, Kruk J, Havaux M, Pospíšil P. 2018. Chemical quenching of singlet oxygen by plastoquinols and their oxidation products in Arabidopsis. Plant Journal 95: 848–861.

Fischer BB, Dayer R, Wiesendanger M, Eggen RIL. 2007. Independent regulation of the GPXH gene expression by primary and secondary effects of high light stress in Chlamydomonas reinhardtii. Physiologia Plantarum 130: 195–206.

Fischer BB, Eggen ÆRIL, Trebst ÆA. 2006. The glutathione peroxidase homologous gene Gpxh in Chlamydomonas reinhardtii is upregulated by singlet oxygen produced in photosystem II. : 583–590.

Fischer BB, Hideg É, Krieger-Liszkay A. 2013. Production, Detection, and Signaling of Singlet Oxygen in Photosynthetic Organisms. Antioxidants & Redox Signaling 18: 2145–2162.

Flors C, Fryer MJ, Waring J, Reeder B, Bechtold U, Mullineaux PM, Nonell S, Wilson MT, Baker NR. 2006. Imaging the production of singlet oxygen in vivo using a new fluorescent sensor, Singlet Oxygen Sensor Green®. Journal of Experimental Botany 57: 1725–1734.

Fufezan C, Gross CM, Sjödin M, Rutherford AW, Krieger-Liszkay A, Kirilovsky D. 2007. Influence of the redox potential of the primary quinone electron acceptor on photoinhibition in photosystem II. Journal of Biological Chemistry 282: 12492–12502.

Funk C, Vermaas W. 1999. A Cyanobacterial Gene Family Coding for Single-Helix Proteins Resembling Part of the Light-Harvesting Proteins from Higher Plants. Biochemistry 38: 9397–9404.

Gao F, Zhao J, Chen L, Battchikova N, Ran Z, Aro EM, Ogawa T, Ma W. 2016. The NDH-1L-PSI supercomplex is important for efficient cyclic electron transport in cyanobacteria. Plant Physiology 172: 1451–1464.

Glaeser J, Nuss AM, Berghoff BA, Klug G. 2011. Singlet Oxygen Stress in Microorganisms. Elsevier Ltd.

Hanaoka M, Tanaka K. 2008. Dynamics of RpaB-promoter interaction during high light stress, revealed by chromatin immunoprecipitation (ChIP) analysis in Synechococcus elongatus PCC 7942. Plant Journal 56: 327–335.

Havaux M, Guedeney G, He Q, Grossman AR. 2003. Elimination of high-light-inducible polypeptides related to eukaryotic chlorophyll a/b-binding proteins results in aberrant photoacclimation in Synechocystis PCC6803. Biochimica et Biophysica Acta - Bioenergetics 1557: 21–33.

Havaux M, Hagemann M, Yeremenko N, Matthijs HCP, Jeanjean R. 2005. The chlorophyll-binding protein IsiA is inducible by high light and protects the cyanobacterium Synechocystis PCC6803 from photooxidative stress. 579: 2289–2293.

He Q, Dolganov N, Björkman O, Grossman AR. 2001. The high light-inducible polypeptides in Synechocystis PCC6803. Expression and function in high light. Journal of Biological Chemistry 276: 306–314.

Hernández-Prieto MA, Semeniuk TA, Giner-Lamia J, Futschik ME. 2016. The transcriptional landscape of the photosynthetic model cyanobacterium Synechocystis sp. PCC6803. Scientific Reports 6: 1–15.

Hideg É, Kós PB, Vass I. 2007. Photosystem II damage induced by chemically generated singlet oxygen in tobacco leaves. Physiologia Plantarum 131: 33–40.

Hideg É, Spetea C, Vass I. 1994. Singlet oxygen production in thylakoid membranes during photoinhibition as detected by EPR spectroscopy. Photosynthesis Research 39: 191–199.

Hihara Y, Kamei A, Kanehisa M, Kaplan A, Ikeuchi M. 2001. DNA microarray analysis of cyanobacterial gene expression during acclimation to high light. Plant Cell 13: 793–806.

Hsiao HY, He Q, Van Waasbergen LG, Grossman AR. 2004. Control of photosynthetic and high-light-responsive genes by the histidine kinase DspA: Negative and positive regulation and interactions between signal transduction pathways. Journal of Bacteriology 186: 3882–3888.

Huckauf J, Nomura C, Forchhammer K, Hagemann M. 2000. Stress responses of Synechocystis sp. strain PCC 6803 mutants impaired in genes encoding putative alternative sigma factors. Microbiology 146: 2877–2889.

Imamura S, Asayama M, Takahashi H, Tanaka K, Takahashi H, Shirai M. 2003. Antagonistic dark/light-induced SigB/SigD, group 2 sigma factors, expression through redox potential and their roles in cyanobacteria. FEBS Letters 554: 357–362.

J. Sambrook, D.W. Russell. 2001. Molecular cloning : a laboratory manual, 3rd ed., Cold Spring Harbor Laboratory Press, Cold Spring Harbor, N.Y.

Järvi S, Suorsa M, Aro EM. 2015. Photosystem II repair in plant chloroplasts - Regulation, assisting proteins and shared components with photosystem II biogenesis. Biochimica et Biophysica Acta - Bioenergetics 1847: 900–909.

Kanesaki Y, Suzuki I, Allakhverdiev SI, Mikami K, Murata N. 2002. Salt stress and hyperosmotic stress regulate the expression of different sets of genes in Synechocystis sp. PCC 6803. Biochemical and Biophysical Research Communications 290: 339–348.

Kappell AD, Bhaya D, Van Waasbergen LG. 2006. Negative control of the high light-inducible hliA gene and implications for the activities of the NblS sensor kinase in the cyanobacterium Synechococcus elongatus strain PCC 7942. Archives of Microbiology 186: 403–413.

Kappell AD, Van Waasbergen LG. 2007. The response regulator RpaB binds the high light regulatory 1 sequence upstream of the high-light-inducible hliB gene from the cyanobacterium Synechocystis PCC 6803. Archives of Microbiology 187: 337–342.

Keun PL, Kim C, Landgraf F, Apel K. 2007. EXECUTER1- and EXECUTER2- dependent transfer of stress-related signals from the plastid to the nucleus of Arabidopsis thaliana. Proceedings of the National Academy of Sciences of the United States of America 104: 10270–10275.

Kirtania P, Hódi B, Mallick I, Vass IZ, Fehér T, Vass I, Kós PB. 2019. A single plasmid based CRISPR interference in Synechocystis 6803 – A proof of concept. PLOS ONE 14: e0225375.

Kodru S, ur Rehman A, Vass I. 2020. Chloramphenicol enhances Photosystem II photodamage in intact cells of the cyanobacterium Synechocystis PCC 6803. Photosynthesis Research 145: 227–235.

Komenda J, Sobotka R. 2016. Cyanobacterial high-light-inducible proteins - Protectors of chlorophyll-protein synthesis and assembly. Biochimica et Biophysica Acta - Bioenergetics 1857: 288–295.

Konert MM, Wysocka A, Koník P, Sobotka R. 2022. High-light-inducible proteins HliA and HliB: pigment binding and protein–protein interactions. Photosynthesis Research.

Krieger-Liszkay A. 2005. Singlet oxKrieger-Liszkay, A. (2005). Singlet oxygen production in photosynthesis. Journal of Experimental Botany, 56(411), 337–346. 10.1093/jxb/erh237ygen production in photosynthesis. Journal of Experimental Botany 56: 337–346.

Krieger-Liszkay A, Fufezan C, Trebst A. 2008. Singlet oxygen production in photosystem II and related protection mechanism. Photosynthesis Research 98: 551–564.

Kufryk G, Hernandez-Prieto MA, Kieselbach T, Miranda H, Vermaas W, Funk C. 2008. Association of small CAB-like proteins (SCPs) of Synechocystis sp. PCC 6803 with Photosystem II. Photosynthesis Research 95: 135–145.

Kunert A, Hagemann M, Erdmann N. 2000. Construction of promoter probe vectors for Synechocystis sp. PCC 6803 using the light-emitting reporter systems Gfp and LuxAB. Journal of Microbiological Methods 41: 185–194.

Ledford HK, Chin BL, Niyogi KK. 2007. Acclimation to Singlet Oxygen Stress in Chlamydomonas reinhardtii L. 6: 919–930.

Leisinger U, Rüfenacht K, Fischer B, Pesaro M, Spengler A, Zehnder AJB, Eggen RIL. 2001. The glutathione peroxidase homologous gene from Chlamydomonas reinhardtii is transcriptionally up-regulated by singlet oxygen. Plant Molecular Biology 46: 395–408.

Lex A, Gehlenborg N, Strobelt H, Vuillemot R, Pfister H. 2014. UpSet: Visualization of intersecting sets. IEEE Transactions on Visualization and Computer Graphics 20: 1983–1992.

Liu J, Qiao J, Shao M, Chen L, Wang J, Wu G, Tian X, Huang S, Zhang W. 2013. Systematic characterization of hypothetical proteins in Synechocystis sp. PCC 6803 reveals proteins functionally relevant to stress responses. Gene 512: 6–15.

Los DA, Zorina A, Sinetova M, Kryazhov S, Mironov K, Zinchenko V V. 2010. Stress sensors and signal transducers in cyanobacteria. Sensors 10: 2386–2415.

Di Mascio P, Martinez GR, Miyamoto S, Ronsein GE, Medeiros MHG, Cadet J. 2019. Singlet Molecular Oxygen Reactions with Nucleic Acids, Lipids, and Proteins. Chemical Reviews 119: 2043–2086.

Michel KP, Pistorius EK. 2004. Adaptation of the photosynthetic electron transport chain in cyanobacteria to iron deficiency: The function of IdiA and IsiA. Physiologia Plantarum 120: 36–50.

Mizusawa N, Tomo T, Satoh K, Miyao M. 2003. Degradation of the D1 Protein of Photosystem II under Illumination in Vivo:L Two Different Pathways Involving Cleavage or Intermolecular Cross-Linking. Biochemistry 42: 10034–10044.

Mohamed A, Eriksson J, Osiewacz HD, Jansson C. 1993. Differential expression of the psbA genes in the cyanobacterium Synechocystis 6803. Molecular and General Genetics MGG 238: 161–168.

Nishiyama Y, Allakhverdiev SI, Yamamoto H, Hayashi H, Murata N. 2004. Singlet Oxygen Inhibits the Repair of Photosystem II by Suppressing the Translation Elongation of the D1 Protein in Synechocystis sp. PCC 6803. Biochemistry 43: 11321–11330.

Nixon PJ, Michoux F, Yu J, Boehm M, Komenda J. 2010. Recent advances in understanding the assembly and repair of photosystem II. Annals of Botany 106: 1–16.

Nuss AM, Adnan F, Weber L, Berghoff BA, Glaeser J, Klug G. 2013. DegS and RseP homologous proteases are involved in singlet oxygen dependent activation of RpoE in Rhodobacter sphaeroides. PLoS ONE 8: 1–15.

Ogawa K, Yoshikawa K, Matsuda F, Toya Y, Shimizu H. 2018. Transcriptome analysis of the cyanobacterium Synechocystis sp. PCC 6803 and mechanisms of photoinhibition tolerance under extreme high light conditions. Journal of Bioscience and Bioengineering 126: 596–602.

Okada K, Ikeuchi M, Yamamoto N, Ono TA, Miyao M. 1996. Selective and specific cleavage of the D1 and D2 proteins of Photosystem II by exposure to singlet oxygen: Factors responsible for the susceptibility to cleavage of the proteins. Biochimica et Biophysica Acta - Bioenergetics 1274: 73–79.

Op Den Camp RGL, Przybyla D, Ochsenbein C, Laloi C, Kim C, Danon A, Wagner D, Hideg É, Göbel C, Feussner I, et al. 2003. Rapid Induction of Distinct Stress Responses after the Release of Singlet Oxygen in Arabidopsis. Plant Cell 15: 2320–2332.

Page MT, McCormac AC, Smith AG, Terry MJ. 2017. Singlet oxygen initiates a plastid signal controlling photosynthetic gene expression. New Phytologist 213: 1168–1180.

Patyi G, Hódi B, Solymosi D, Vass I, Kós PB. 2021. Increased sensitivity of heavy metal bioreporters in transporter deficient Synechocystis PCC6803 mutants. Plos One 16: e0261135.

Peca L, Kós PB, Máté Z, Farsang A, Vass I. 2008. Construction of bioluminescent cyanobacterial reporter strains for detection of nickel, cobalt and zinc. FEMS Microbiology Letters 289: 258–264.

Peca L, Kós PB, Vass I. 2007. Characterization of the activity of heavy metal-responsive promoters in the cyanobacterium Synechocystis PCC 6803. Acta Biologica Hungarica 58: 11–22.

Peca L, Nagy CI, Ôrdog A, Vass I, Kós PB. 2017. Development of a bioluminescent cyanobacterial reporter strain for detection of arsenite, arsenate and antimonite. Environmental Engineering and Management Journal 16: 2443–2450.

Pospíšil P, Kumar A, Prasad A. 2022. Reactive oxygen species in photosystem II: relevance for oxidative signaling. Photosynthesis Research 152: 245–260.

Prasad A, Ferretti U, Sedlaová M, Pospíšil P. 2016. Singlet oxygen production in Chlamydomonas reinhardtii under heat stress. Scientific Reports 6: 1–13.

Promnares K, Komenda J, Bumba L, Nebesarova J, Vacha F, Tichy M. 2006. Cyanobacterial small chlorophyll-binding protein ScpD (HliB) is located on the periphery of photosystem II in the vicinity of PsbH and CP47 subunits. Journal of Biological Chemistry 281: 32705–32713.

Ramel F, Birtic S, Ginies C, Soubigou-taconnat L, Triantaphylidès C. 2012. Carotenoid oxidation products are stress signals that mediate gene responses to singlet oxygen in plants. 109.

Rehman AU, Cser K, Sass L, Vass I. 2013. Characterization of singlet oxygen production and its involvement in photodamage of Photosystem II in the cyanobacterium Synechocystis PCC 6803 by histidine-mediated chemical trapping. Biochimica et Biophysica Acta - Bioenergetics 1827: 689–698.

Riethman H, Bullerjahn G, Reddy KJ, Sherman LA. 1988. Regulation of cyanobacterial pigment-protein composition and organization by environmental factors. Photosynthesis research 18: 133–161.

Rippka R, Deruelles J, Waterbury JB. 1979. Generic assignments, strain histories and properties of pure cultures of cyanobacteria. Journal of General Microbiology 111: 1–61.

Shao N, Duan GY, Bock R, Molekulare M. 2013. A Mediator of Singlet Oxygen Responses in Chlamydomonas reinhardtii and Arabidopsis Identi fi ed by a Luciferase-Based Genetic Screen in Algal Cells. 25: 4209–4226.

Shoumskaya MA, Paithoonrangsarid K, Kanesaki Y, Los DA, Zinchenko V V., Tanticharoen M, Suzuki I, Murata N. 2005. Identical Hik-Rre systems are involved in perception and transduction of salt signals and hyperosmotic signals but regulate the expression of individual genes to different extents in Synechocystis. Journal of Biological Chemistry 280: 21531–21538.

Shumbe L, Alessandro SD, Shao N, Chevalier A, Ksas B, Bock R, Havaux M, Cadarache CEA, Umr C, Université A, et al. 2017. Original Article METHYLENE BLUE SENSITIVITY 1 (MBS1) is required for acclimation of Arabidopsis to singlet oxygen and acts downstream of β -cyclocitral. 1: 216–226.

Silva P, Thompson E, Bailey S, Kruse O, Mullineaux CW, Robinson C, Mann NH, Nixon PJ. 2003. FtsH is involved in the early stages of repair of photosystem II in Synechocystis sp PCC 6803. The Plant cell 15: 2152–2164.

Sinha RK, Komenda J, Knoppová J, Sedlářová M, Pospíšil P. 2012. Small CAB-like proteins prevent formation of singlet oxygen in the damaged photosystem II complex of the cyanobacterium Synechocystis sp. PCC 6803. Plant, Cell and Environment 35: 806–818.

Strand A, Asami T, Ecker JR. 2003. Chloroplast comunication triggered by accumulation of Mg-protoporphyrinknix. Nature 421: 79–83.

Tomo T, Kusakabe H, Nagao R, Ito H, Tanaka A, Akimoto S, Mimuro M, Okazaki S. 2012. Luminescence of singlet oxygen in photosystem II complexes isolated from cyanobacterium Synechocystis sp. PCC6803 containing monovinyl or divinyl chlorophyll a. Biochimica et Biophysica Acta - Bioenergetics 1817: 1299–1305.

Turunen O, Koskinen S, Kurkela J, Karhuvaara O, Hakkila K, Tyystjärvi T. 2022. Roles of Close Homologues SigB and SigD in Heat and High Light Acclimation of the Cyanobacterium Synechocystis sp. PCC 6803. Life 12.

Vass I. 2012. Biochimica et Biophysica Acta Molecular mechanisms of photodamage in the Photosystem II complex IZ. BBA - Bioenergetics 1817: 209–217.

Vass I, Cser K. 2009. Janus-faced charge recombinations in photosystem II photoinhibition. Trends in Plant Science 14: 200–205.

Wang L, Kim C, Xu X, Piskurewicz U, Dogra V, Singh S, Mahler H, Apel K. 2016. Singlet oxygen- and EXECUTER1-mediated signaling is initiated in grana margins and depends on the protease FtsH2. Proceedings of the National Academy of Sciences of the United States of America 113: E3792–E3800.

Zavafer A. 2021. A theoretical framework of the hybrid mechanism of photosystem II photodamage. Photosynthesis Research 149: 107–120.

